# Overlap across psychotic disorders: A functional network connectivity analysis

**DOI:** 10.1101/2022.03.13.484190

**Authors:** Hossein Dini, Luis E. Bruni, Thomas Z. Ramsøy, Vince D. Calhoun, Mohammad S. E. Sendi

## Abstract

Functional network connectivity has previously been shown to distinguish patient groups from healthy controls (HC). However, the overlap across schizophrenia (SZ), bipolar (BP), and schizoaffective disorder (SAD) is not clear yet. This study focuses on finding overlap across these three psychotic disorders using dynamic FNC (dFNC) and compares it with static FNC (sFNC). We used resting-state fMRI, demographics, and clinical information from the Bipolar– Schizophrenia Network on Intermediate Phenotypes cohort. The data includes three groups of patients with schizophrenia (SZP, N=102), bipolar (BPP, N=102), and schizoaffective (SADP, N=102), their relatives SZR (N=102), BPR (N=102), SADR (N=102), and HC (N=118) groups. After estimating each individual’s dFNC, we put them into three identical states. We estimated five different features, including occupancy rate (OCR), number of transitions, the total number of transitions, and the total distance traveled. Finally, the extracted features are tested statistically across patients and HC groups. In addition, we explored the link between the clinical scores and the extracted features. We found that the OCR difference between SZP and SZR in state2, between BPP and HC in state1, and between SADP and HC in state2. Also, state2 OCR separates SZP from BPP, state 3 OCR separates BPP from SZP and SADP. Moreover, the OCR and traveled distance feature extracted from SZ and BP could significantly predict PANSS Total and PANSS General scores. Finally, combined distance features of all disorders showed a significant relationship to PANSS Total and PANSS General scores.

## 1. Introduction

Previous psychiatric disorder studies have found that schizophrenia (SZ) and bipolar (BP) disorders show some identical symptoms such as delusions, hallucinations, and mood disturbance (Pearlson, 2015). It has been argued that BP has a relatively high misdiagnosis rate, and its diagnosis can be mixed with SZ (Mukherjee et al., 1983). Rokham et al. provided evidences showing the traditional diagnostic labels are not necessarily valid by implementing a machine learning algorithm to detect the noisy label on a data including SZ, BP, and SAD disorders (Rokham et al., 2020). Meyer et al. mentioned that about 45% of people suffering from bipolar are misdiagnosed with SZ due to having hallucinations (Meyer and Meyer, 2009). Also, Shen et al. reported that 20%-30% of BP patients are diagnosed as SZ mistakenly (Shen et al., 2018). Additionally, recent studies argued that schizoaffective disorder (SAD) is placed somewhere between SZ and BP spectrum (Tamminga et al., 2014; Mancuso et al., 2015; Paudel et al., 2020). SAD was first defined by Kasanin as having combined symptoms of SZ and mood disorder (Kasanin, 1933). Also, the diagnostic and statistical manual for mental disorders, fifth edition (DSM-V) considers SAD as an independent diagnosis under SZ spectrum disorders, acknowledging their symptomatic overlap (Association and Association, 2013).

In addition to clinical overlap in SZ, BP, and SAD (Laursen et al., 2009; Cosgrove and Suppes, 2013; Malaspina et al., 2013; Du et al., 2020b), some neuroimaging studies have also reported the existence of overlap among three disorders (Ivleva et al., 2013; Amann et al., 2016; Birur et al., 2017). Ivleva et al. showed that SZ and SAD have a similar amount of gray matter reduction in many brain regions while BP has less gray matter reduction, which is limited to the frontotemporal cortex (Ivleva et al., 2013). In a voxel-based morphometry study, Amann et al. compared three disorders with a healthy (HC) group and concluded that SAD and SZ have similar volume reduction in overlapping areas together, where this reduction had less similar patterns to than BP (Amann et al., 2016). A review paper by Birur et al. reported that SZ and BP have comparable and consistent white matter impairment, while gray matter reduction is higher in SZ than in BP (Birur et al., 2017). Finally, Yan et al. reported the existence of relation across SZ, BP, and SAD disorders implementing a deep classification and clustering framework on extracted features from correlation between different brain regions (Yan et al., 2021).

Psychiatric disorders are typically diagnosed based on clinical symptoms and DSM-V criteria. Nowadays, there is an emerging trend towards the diagnosis of such disorders based on electrophysiological biomarkers (Singh and Rose, 2009). Identifying electrophysiological biomarkers as an alternative for behavioral signs and symptoms, which leads to subjective diagnosis, might provide a more precise and reliable diagnosis (Moffitt et al., 2008). Moreover, biomarkers can sub-categorize disorders based on physiological criteria and provide a personalized approach to psychiatric treatments (Yan et al., 2021). In addition, biomarkers can be considered not only as a diagnostic approach but also as an assist to tailor treatments. However, it is mentioned that defining biomarkers using current methods is challenging (Singh and Rose, 2009). As genetic, molecular, histological, and neuroimaging studies suggest, major psychotic disorders such as SZ, BP, and SAD share overlapping features (Tamminga et al., 2013, 2014; Clementz et al., 2020; Yan et al., 2021), but to the best of our knowledge, the degree of overlap in their neuroimaging features is not well established.

Identifying biomarkers from brain functional connectivity derived from functional magnetic resonance imaging (fMRI) data is a potential supportive/alternative measure for symptom-based clinical features to diagnose/modify treatment for psychiatric disorders (Sendi et al., 2021c)(Stephan et al., 2017; Du et al., 2018b, 2020b). Recently, functional network connectivity (FNC), which is obtained evaluating the temporal dependence over the entire resting-state fMRI (rs-fMRI) time series (called static functional network connectivity analysis), has been demonstrated to be highly informative to reveal underlying brain connectivity patterns in mental disorders (Mulders et al., 2015; Yan et al., 2019; Dini et al., 2020; Liu et al., 2020; Luo et al., 2021). Previous studies used various features including spatial functional networks, and FNC based on independent component analysis (ICA) to discriminate BPP and SZ from the HC group (Arribas et al., 2010; Dini et al., 2020),(Jafri et al., 2008). Xia et al., in a study investigating functional network features (e.g., clustering coefficient) of BP, SZ, and major depressive disorder, reported that their network features go toward randomized configurations with different degrees. Since SAD patients are usually categorized in the same group as SZ (Du et al., 2020b), fewer studies have investigated the group overlap in group discriminative neurophysiological features across SZ, BP, and SAD simultaneously. One study evaluated the spatial brain network of all three diseases. They implemented an approach called group information-guided ICA method on the data and the results were able to discriminate the disorders (Du et al., 2015b). However, further investigation is needed to understand the underlying neural mechanism and specify features across these three disorders (Colibazzi, 2014).

The estimated FNC in the abovementioned studies is assumed to be static, but this assumption does not account for the dynamic nature of brain FNC. Dynamic FNC (dFNC) was introduced to overcome this limitation (Allen et al., 2014; Zendehrouh et al., 2020; Sendi et al., 2021f). dFNC estimates FNC in shorter time intervals of time series compared to static FNC, which averages over the entire dataset (Calhoun et al., 2014). In recent years, dFNC has been found to be more sensitive and informative than static FNC in underlying various brain regions connectivity of different mental disorders such as SZ, major depressive disorder, and Alzheimer’s disease (Rashid et al., 2016; Du et al., 2017; Sendi et al., 2020, 2021f, 2021a). Previous studies reported abnormalities in dFNC of SZ compared to the HC group population, and different state activity between SZ and BP (Damaraju et al., 2014; Du et al., 2016). Du et al. decomposed connectivity patterns of SZ and HC group to different states and found that participants with high psychosis risk show intermediate pattern between HC and SZ group (Du et al., 2018a). Finally, investigation of the dFNC of BP, SZ, and SAD suggested both hypoconnectivity and hyperconnectivity were present from HC to BP to SAD to SZ, in a decreasing and increasing trend, respectively. They also reported that dynamic FNC revealed more detailed group differences than static FNC.

The current study focuses on finding group discriminative features that modulate clinical overlap between psychotic disorders (i.e., SZ, BP, and SAD), predicting the clinical outcome. We first calculate dFNC via a sliding window approach and then used k-means clustering to estimate three different states to see how the brain activity alters across different states. Finally, we calculated various features from the activity of extracted states and explored overlap within (e.g., SZ versus HC, BP versus HC, and SAD versus HC) and between (e.g., SZ, BP, and SAD) disorders to identify group discriminating features that are sensitive to overlap. Next, we investigate the possibility of predicting clinical data using identified group discriminative features. Finally, we calculate static FNC to compare it with the dFNC results.

## 2. Materials and Methods

### 2.1. Participants and Clinical Metrics

This study used neuroimaging, clinical, and demographic information of 730 participants in total from Bipolar and Schizophrenia Network on Intermediate Phenotypes project (Moffitt et al., 2008). This dataset includes patient with schizophrenia (SZP), bipolar (BPP), schizoaffective disorder (SADP), their relative (i.e., SZR, BPR, and SADR), and HC. The demographic and clinical information of the participants is provided in Table 1. for the exclusion criteria was: 1) having any other neurodegenerative and neurological disorders other than the abovementioned disorders, 2) having an addiction to alcohol or drugs, and pregnancy, and 3) being susceptible to danger from the MRI scanner, such as using a non MRI-compatible pacemaker(Dini et al., 2021).

**Table 1:** clinical and demographical information for different groups

|  | SZP | BPP | SADP | SZR | BPR | SADR | HC |
| --- | --- | --- | --- | --- | --- | --- | --- |
| *Number*<br>*(Actual/Chosen)* | 102 | 102 | 102 | 102 | 102 | 102 | 118 |
| *Age* | M:34.34<br>SD:12.49 | M:34.71<br>SD:12.35 | M:35.17<br>SD:12.65 | M:34.45<br>SD:14.16 | M:34.44<br>SD:14.50 | M:34.05<br>SD:14.36 | M:33.75<br>SD:12.77 |
| *Gender(M/F)* | 47/55 | 34/68 | 46/56 | 36/66 | 43/59 | 34/68 | 44/74 |
| *PANSS Positive* | M:16.62<br>SD:5.77 | 13.04<br>4.56 | 18.04<br>5.11 | N.A | N.A | N.A | N.A |
| *PANSS Negative* | M:16.28<br>SD:5.57 | M:12.04<br>SD:3.92 | M:15.39<br>SD:4.34 | N.A | N.A | N.A | N.A |
| *PANSS Gen-total* | M:31.88<br>SD:8.32 | M:29.05<br>SD:8.25 | M:34.58<br>SD:9.12 | N.A | N.A | N.A | N.A |
| *PANSS total* | M:64.79<br>SD:16.48 | M:54.13<br>SD:14.03 | M:67.93<br>SD:15.79 | N.A | N.A | N.A | N.A |

### 2.2. Data Acquisition and Pre-processing

The data was collected from participants in the multi-site Bipolar and Schizophrenia Network on Intermediate Phenotypes project (Tamminga et al., 2013; Meda et al., 2014, 2015). The fMRI scans occurred at six different sites (Baltimore, Boston, Chicago, Dallas, Detroit, and Hartford). For all sites, the scanning took roughly five minutes. At the time of the study, all of the participants were psychiatrically stable and on stable drug regimens. Participants were told to close their eyes and stay awake. The detailed information of the sites can be seen in Table S13. All the collected data went through the standard preprocessing steps of fMRI data, implementing statistical parametric mapping (SPM12, https://www.fil.ion.ucl.ac.uk/spm/) in the MATLAB 2021 environment. We removed the first five fMRI scans to remove the noises generated from participants’ adaptation to the scanner and longitudinal relaxation effects. Then, we corrected the temporal differences in slice acquisition, and head and body motions were corrected using the SPM toolbox and rigid body motion correction, respectively. Next, we spatially normalized all the imaging data using echo-planar imaging (EPI) template in standard Montreal Neurological Institute (MNI) space and the resampled to 3 ×3 ×3 *mm*^3^. Finally, we applied a 6-mm full-width half-maximum (FWHM) Gaussian kernel on the data to smooth it spatially.

We extracted reliable independent components (ICs), including Subcortical Network (SCN), Auditory Network (ADN), Sensory Motor Network (SMN), Visual Network (VSN), Cognitive Control Network (CCN), Default Mode Network (DMN), and cerebellum (CBN). To this aim, we implemented the Neuromark fully automated group ICA pipeline Figure1A, using GIFT (http://trendscenter.org/software/gift). This method uses previously extracted components as priors for spatially constrained ICA. In this method, replicable ICs are identified and used for spatial priors, using group-level spatial maps from two large-sample HC datasets (Du et al., 2020a).

Before applying the dynamic functional connectivity method, we used the following steps to reject the artifacts and denoise the data; first, linear, quadratic, and cubic de-trending is implemented. Then, we used six realignment parameters and their temporal derivatives for multiple regression. Finally, we removed outliers and filtered the data using a bandpass filter with 0.01 to 0.15 Hz cutoff frequencies. The list of identified components in the abovementioned regions can be seen in Table 2.

**Table 2:**
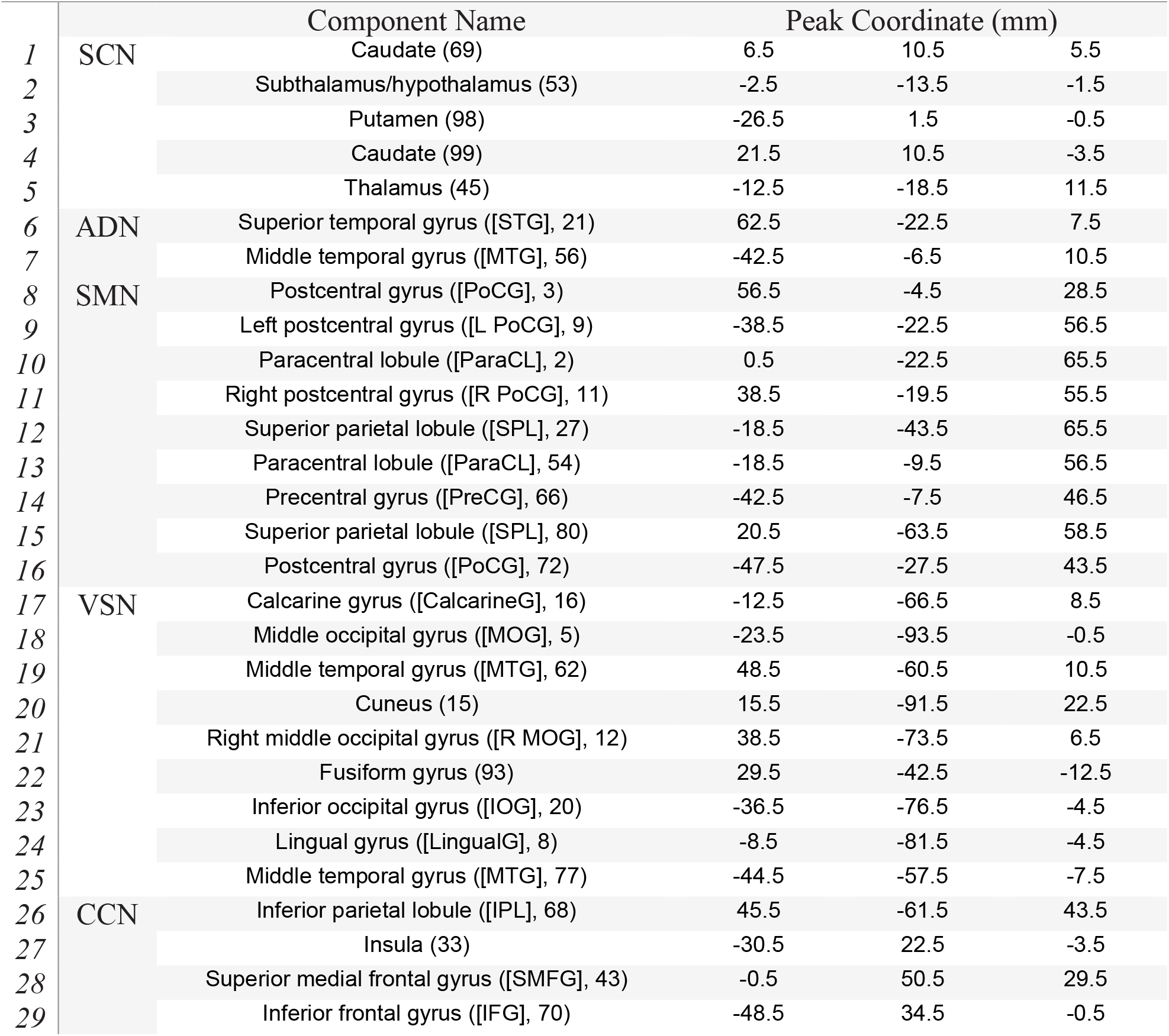
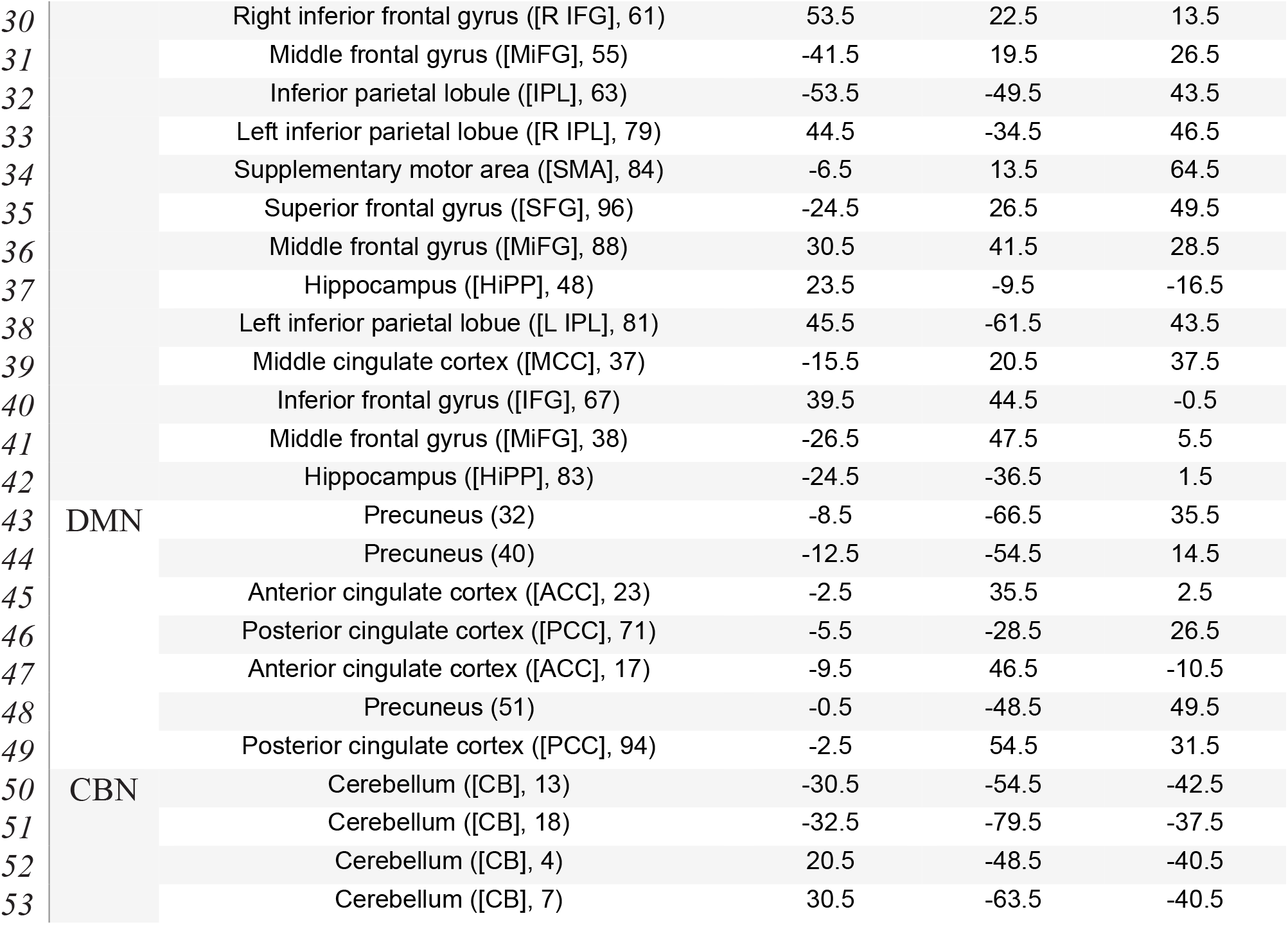
Component labels

### 2.3. Functional Network Connectivity

To calculate dynamic functional network connectivity (dFNC), we defined a window by convolving a rectangle (window size = 20 TRs = 40 s) with a Gaussian (σ = 3 s) for localizing each time point in the dataset. Then, by sliding this window through the data, we calculated the Pearson correlation coefficient between 53 nodes from defined regions within each window Figure1B. With 53 subnodes, we obtained 1378 identical connectivity features from each window. Then we concatenated extracted features of participants in a *C* ×*C* ×*T* array (where C=53 denotes the number of ICs, and T= 178 denotes the number of windows). Finally, we concatenated the participant’s calculated arrays to show brain connectivity changes between defined regions as a function of time as shown in Figure1C (Allen et al., 2014; Dini et al., 2021; Sendi et al., 2021d, 2021b). Moreover, we calculated static FNC of each subject by calculating the Pearson correlation across all 53 regions for the entire rs-fMRI session (Figure1B). In total, we obtained 1378 identical connectivity features for each subject.

### 2.4. Clustering and dFNC Latent Features

To divide the data into different clusters, we used a k-means clustering algorithm (Figure1D) with the city-block distance method on the output of the previous step, which is concatenated dFNC of all subjects (Allen et al., 2014; Sendi et al., 2021f). Then, we used the Calinski-Harabasz clustering evaluation criterion to find the optimum number of clusters, which is a common method to find optimum numbers of clusters and it fits well with the k-means clustering method (Damaraju et al., 2014; Sendi et al., 2021f). It divides within-cluster distance by between cluster distance and defines a ratio out of that, and uses this ratio as the objective function to minimize it. As the output, we found that three is the optimum number of clusters in this data by searching from K=2 to 9 (Sendi et al., 2020). Furthermore, applying the city block distance method with 100 iterations, we found three distinct states for the group of all participants and a state vector for each individual (Figure1E). This state vector shows how the brain network changes between any pairs of extracted clusters. Out of this state vector, we extracted five identical features, including occupancy rate (OCR), hidden Markov model (HMM), Number of transitions to a state, all number of transitions, and traveled distance (Figure1F).

**Figure 1:**
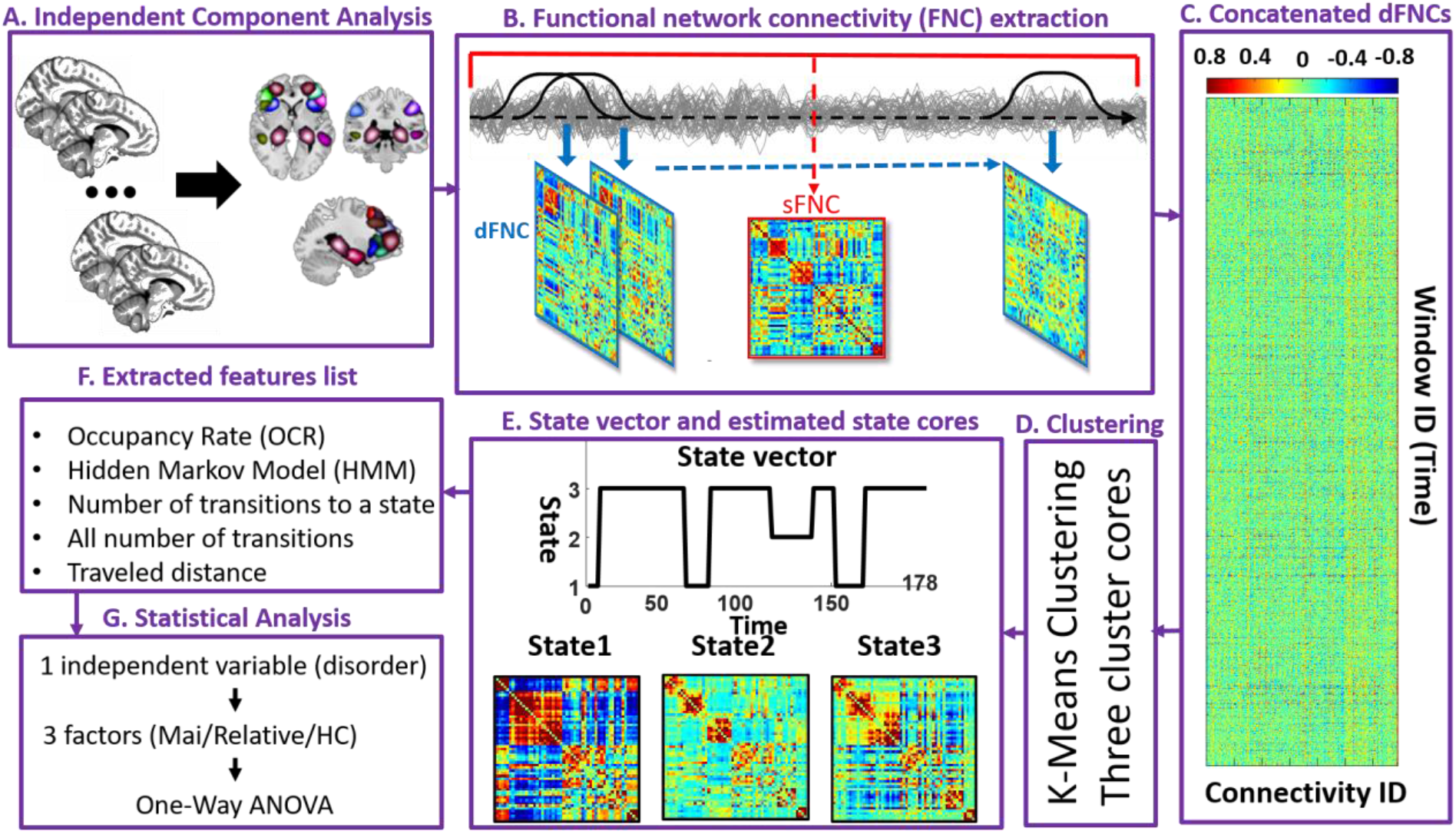
Analytic pipeline: **a**. Using group independent component analysis (ICA), the time-course signal of 53 components of the whole brain has been identified. **b**. dFNC and sFNC calculation. For dFNC, taper sliding window was used to segment the time-course signals and then calculated the functional network connectivity (FNC) of each window. Matrices with blue border indicates the dFNC. For sFNC we applied a big window (indicated with red) on the whole signal. The red bordered matrix indicates sFNC. **c**. All calculated FNCs are concatenated together and are fed to k-mean (k=3) clustering function. Elbow criteria were used to find the optimal k. **d**. Applying clustering method resulted in three identical states and a state vector for each individual (here is an example of state vector for 15^th^ subject). e. Clustering method **f**. based on the state vector of each subject, total of five features were calculated from the state vector of each subject. **g**. We compared the extracted features within and between disease, where both categories has three factors, using one-way ANOVA. We adjusted all p values by the Benjamini-Hochberg false discovery rate (FDR) correction in each analysis.

The OCR is calculated based on the state vector of each participant. This feature shows the proportional time interval that each subject spent in each of the three states. Therefore, we will have three OCR features for each participant. To calculate HMM feature, we estimated the probability of changing from a specific state (e.g., A) to the same (A) or any other existing states (e.g., B). The probability is calculated by calculating the number of transitions from state A to state B and dividing it by the all number of transitions. Since we have three states, all possible probabilities are 9, and therefore, we have 9 HMM features for each participant. To calculate number of transitions to a state, we considered any transition from any state to the target state as one travel. Considering three states, we will have three transitions to a state features per participant. all number of transition is obtained by summing up the calculated number of transitions per state. To determine the traveled distance feature, we used the Euclidian distance method. To this aim, we calculated the distance between any subsequent windows of dFNC matrix and summed up all the distances. This feature is not state-specific, so we have one traveled distance for each subject.

### 2.5. Statistical Analysis

The calculated features are tested statistically between and within disorders (Figure1G). We have one independent variable with three factors (in within disorder the factors are patient group, the relative group, and HC group, and in between disorders the factors are three different disorders). Therefore, we used one-way ANOVA to test the differences, and then we corrected p-values based on the number of statistical test repetition (FDR correction) using the Benjamini-Hochberg method (Gerstung et al., 2014). Moreover, to see whether there is a link between dFNC activity and behavioral scores, we did a partial correlation analysis between calculated features and all four PANSS scores using partial correlation analysis with having age, gender, and TR as covariates.

## 3. Results

This section provides the obtained results from dFNC analysis and compares them in different groups. It comprised clinical results, dynamic and static FNC comparison, dFNC states that are resulted from clustering analysis, statistical results from the comparison of extracted features in different conditions, and the correlation analysis results using five extracted features and four clinical scores.

### 3.1. Clinical outcomes

The clinical and demographical information of all groups is shown in Table 1. The clinical information is comprised of four reports collected from participants except for relatives and healthy controls. Age and gender did not show significant differences among all possible combinations of patient, relative, and HC groups. Table 3 shows the results of the statistical comparison (using two samples t-test, considering the normality of the data) between all possible combinations of the disorders in patient group.

**Table 3:** P-values resulted from comparing all combinations of groups and scores

|  | SZP vs BPP | SZP vs SADP | BPP vs SADP |
| --- | --- | --- | --- |
| PANSS Positive | p=9.51*10<sup>-4</sup> | p=0.22 | p=6.80*10<sup>-8</sup> |
| PANSS Negative | p=2.17*10<sup>-5</sup> | p=0.71 | p=2.04*10<sup>-4</sup> |
| PANSS Gen-total | p=0.09 | p=0.18 | p=5.34*10<sup>-4</sup> |
| PANSS total | p=2.93*10<sup>-4</sup> | p=0.33 | p=5.22*10<sup>-5</sup> |

### 3.2. Dynamic functional network connectivity States

We found three as the optimum number of clusters for the current data is three using Calinski-Harabasz clustering evaluation criterion (Damaraju et al., 2014; Sendi et al., 2021f). Then by applying the k-means clustering method on the data, we discriminated the data into three distinct clusters (states). The clustering evaluation criterion and k-means method are applied on combined dFNC of all selected participants of all different groups and all regions of the brain. Since we aim to compare the overlap between different disorders, we will focus on the whole-brain networks, not looking at the different brain networks separately. The activity of these states is shown in Figure 2. Although this study aims not to evaluate different brain regions individually, it is worthy of mentioning how different regions are active, corresponding to different states, to see the highlighted characteristics of each state. As Figure 2 shows, the connectivity of all regions (both within a specific region and connectivity of a region with others) is higher in state 1 than the other two states. The highest connectivity is within SCN, VSN, and CBN networks, in all three states. The other pronounced difference is between state 2 and state 3 where the correlation of DMN and CCN with all other regions is higher in state 3.

**Figure 2:**
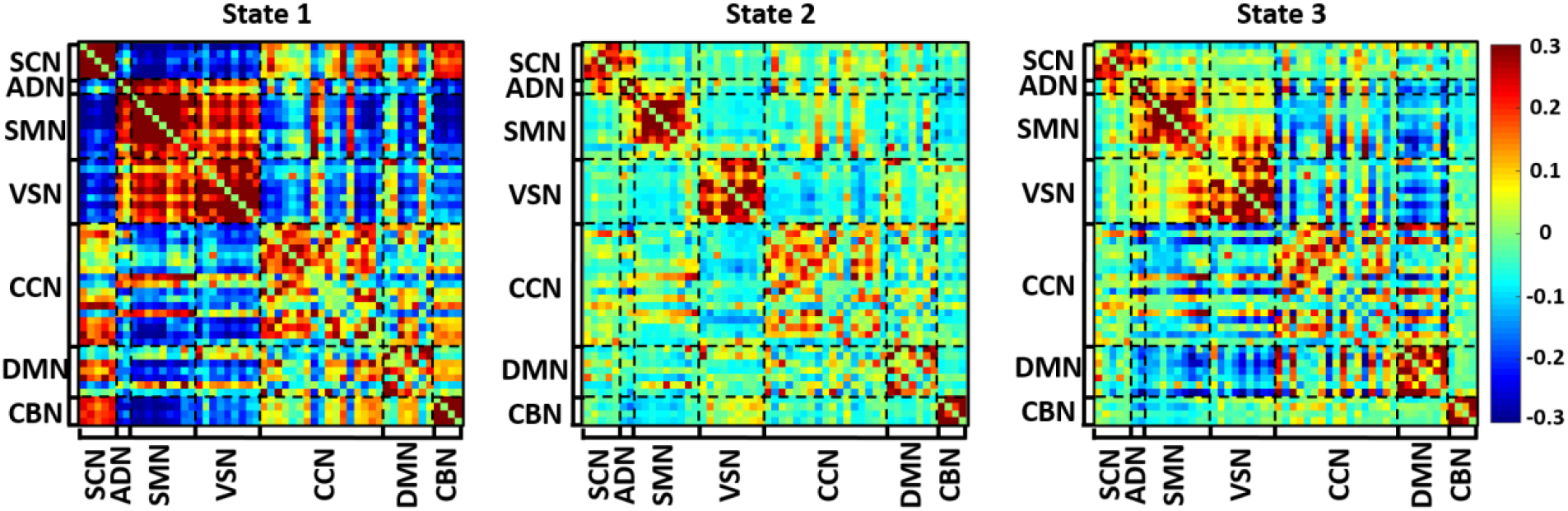
Obtained three states from implementing clustering method on dFNC. Each state is divided to seven regions which are: Subcortical Network (SCN), Auditory Network (ADN), Sensory Motor Network (SMN), Visual Network (VSN), Cognitive Control Network (CCN), Default Mode Network (DMN), and Cerebellum Network(CBN).

### 3.3. Comparison of extracted features within and between group

As mentioned in the method part, we extracted five features: OCR, HMM, number of transitions to a state, number of transitions, and traveled distance. In addition, we divided the patient group into two categories: within disorder (i.e., within SZ, within BP, within SAD vs HC) and between disorder (i.e., between patients across three disorders). In within disorder comparison, we compared the patients, relative, and HC groups together in each state using one-way ANOVA. In between disorder comparison, we compared the values of different disorder together (e.g., SZP, BPP, and SADP) using the same statistical procedure. This paper will elaborate on the significant differences that we found, but the complete information of comparisons is provided in Table S1 to Table S4. Figure 3 shows the statistical comparisons using the OCR feature. Figure 3A to Figure 3C show the within disorder comparison, and Figure 3D and Figure 3E show the between disorder comparison. Figure 3A, shows the differences within SZ where there is 1) a significant difference between patient and relative group (corrected *p*=0.021); 2) the relative and HC group (corrected *p*< 0.001), both in state 2. Figure 3B shows the differences within BP where there is a significant difference between patient and HC group (corrected *p*= 0.045) in state 1. Figure 3C shows the differences within SAD where there is a significant difference between patient and HC group (corrected *p*= 0.001) in state 2. Figure 3D shows the differences between disorders, where the OCR of the SZP group is significantly higher than the BPP one (corrected *p*= 0.012) in state 2 but substantially lower than BPP (corrected *p*= 0.004) in state 3. Moreover, it shows that BPP has significantly higher OCR than SADP (corrected *p*=0.017) in state 3. Finally, Figure 3E shows no significant differences across relative in disorders in none of the states. It is worthy of mentioning that the trend of OCR values in state 1 and state 2 is identical in the within disorder comparison. In Figure 3A to Figure 3C, the OCR of the HC group is higher than relative, and that is higher than the patients (e.g., HC>SZR>SZP) both in state 1 and state 2 of all different disorders, showing an overlap in dFNC features of different disorders. In state 3, the same pattern was observed for SZ and SAD but not for BP disorder. Moreover, in the between disorder analysis (Figure 3D and Figure 3E), it is shown that the OCR of BPP is lower than two other disorders in state 2 and it is higher than other disorders in state 3, and this happened for both patients and relatives.

**Figure 3:**
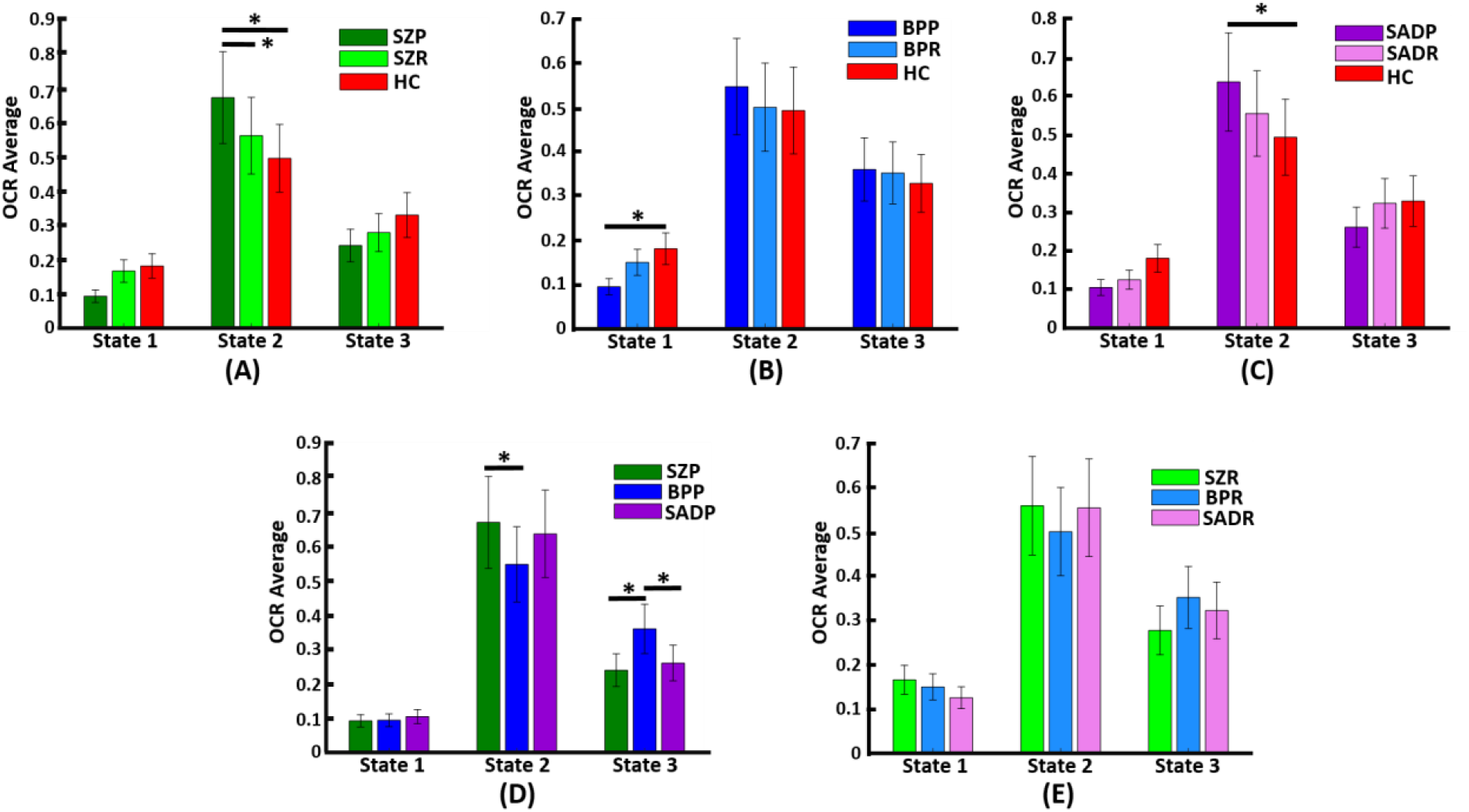
OCR values for three different state for both within and between disease comparison. Panel A to C are related to within diseases comparison, and show OCR values for SZ, BP, and SAD disease, respectively. In panel A, we used dark green color for SZP, light green for SZR and red for HC. It shows there is a significant difference between SZP and SZR (corrected p=0.021) and between SZP and HC (corrected p<0.001.) in state2. In panel B, dark blue shows BPP and light blue shows BPR, and it indicates a significant difference between BPP and HC group (corrected p=0.045) in state1. In panel C dark and light purple are related to SADP and SADR, respectively. It shows a significant difference between SADP and HC group (corrected *p*= 0.001) in state 2. Panel D and E shows average OCR of between disease and are related to main diseases and relative disease, respectively. In panel D, there is a significant difference between SZP and BPP in state2 (corrected p= 0.012), a significant difference between SZP and BPP in state 3 (corrected *p*= 0.004), and a significant difference between BPP and SADP in state 3 (corrected *p*= 0.017). Panel E shows there is not any difference between relative diseases.

There is a significant difference between the within SZ and within the SAD analysis for the HMM feature. Within SZ, the HMM value of SZP is significantly lower than the HC group (corrected p= 0.019) in staying in state 1 (transition from state 1 to state 1). Moreover, in transition from state 3 to state 1, the SZP values are significantly lower than the HC group (corrected p= 0.016). Within SAD, staying in state 2 (transition from state 2 to state 2) is significantly higher in SADP than HC group (corrected p= 0.020). No other significant differences were found in neither HMM values for within BP and between disorders comparison.

We neither found any significant differences within the disorder nor in-between disorder analysis regarding the other features. The only exception was within SZ of the number of transitions to a state feature, which happened in state 2. The value of SZP is significantly lower than the HC group (corrected *p*= 0.029). The detailed information of within/between disorder comparison is provided in Table S1 to Table S4.

### 3.4. Dynamic FNC features links with clinical outcome

We ran a partial correlation method considering age, gender, TR, and biotype covariates. Such analysis aims to investigate whether each extracted feature patient group (i.e., SZP, BPP, and SADP) can predict each clinical score (i.e., PANSS Negative, PANSS Positive, PANSS General, and PANSS Total). Therefore, we used all the computed features in each disorder separately as input, and the significant results are shown in Figure 4, and the rest of the results can be seen in Table S5 to Table S8. Figure 4A shows that the OCR extracted from the BPP in state 2 showed a significant relationship to the general PANSS (*r*=0.269, *p*=0.043). Figure 4B and Figure 4C shows that the OCR and distance feature extracted from SZP has significant link to general PANSS (*r*= 0.257, *p*= 0.015) and total PANSS (*r*=0.233, *p*=0.027), respectively. Finally, Figure 4D highlights that the traveled distance extracted from BPP significantly relates to PANSS General (*r*=0.243, *p*=0.027). In addition to evaluating the predictability of clinical scores in each patient group separately, we tried to predict the scores by combining all of the patients in each extracted feature. Figure 5A and Figure 5B shows that combined distance features of all patients can predict general PANSS (*r*= 0.159, *p*=0.009) and total PANSS (*r*=0.143, *p*=0.020), respectively. The details of the results can be seen in Table S9 to Table S12.

**Figure 4:**
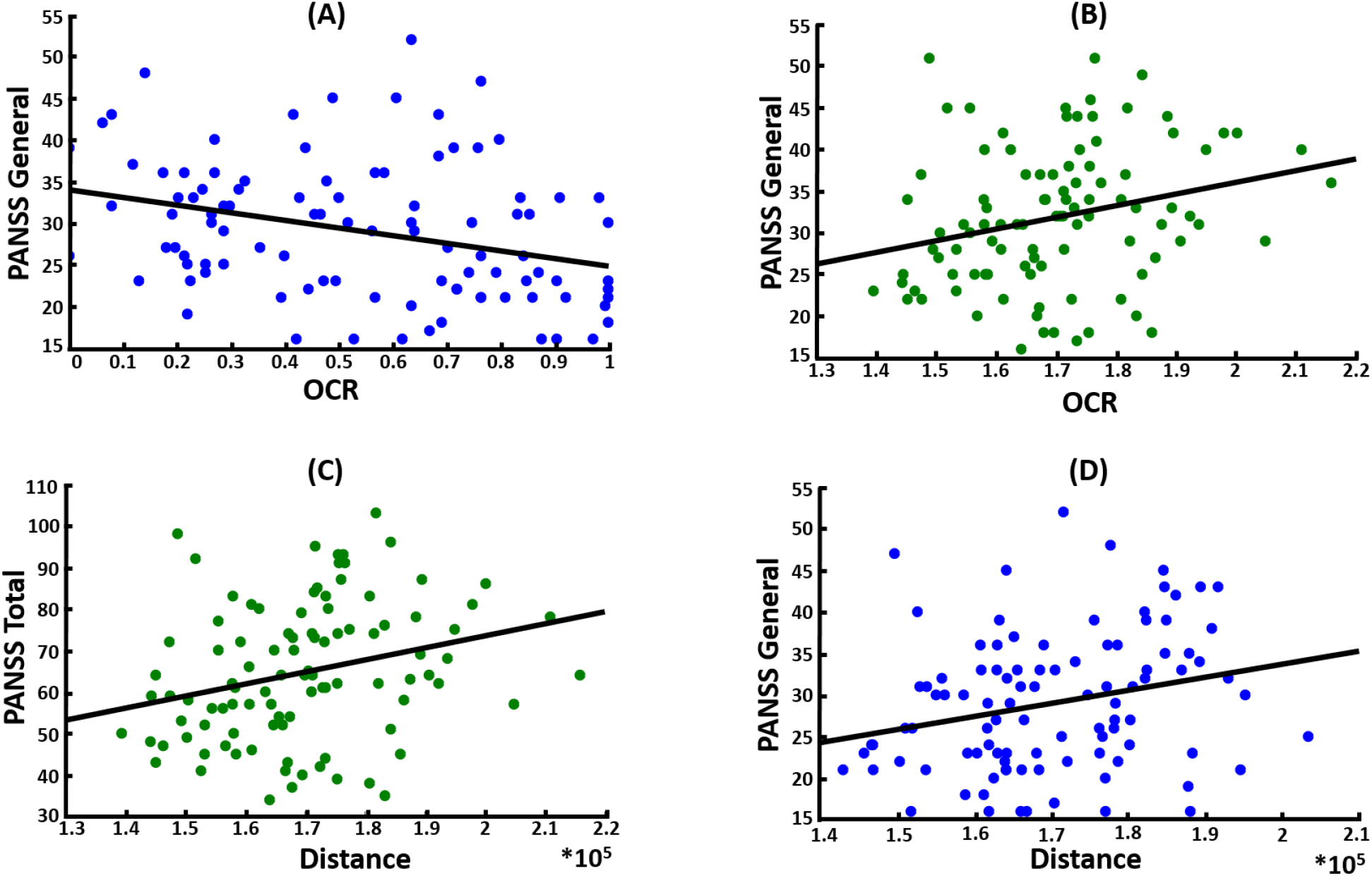
shows correlation analysis for main diseases, considering each disease separately. Panel A shows that OCR is significantly predicting PANSS General scores in BPP (*r*=0.269, *p*=0.043). Panel B is the same as panel B but for SZP (*r*= 0.257, *p*= 0.015). Panel C indicates that distance feature can significantly predict PANSS total in SZP (*r*=0.233, *p*=0.027). Panel D is related to prediction of PANSS General with distance feature in BPP (*r*=0.243, *p*=0.027).

**Figure 5:**
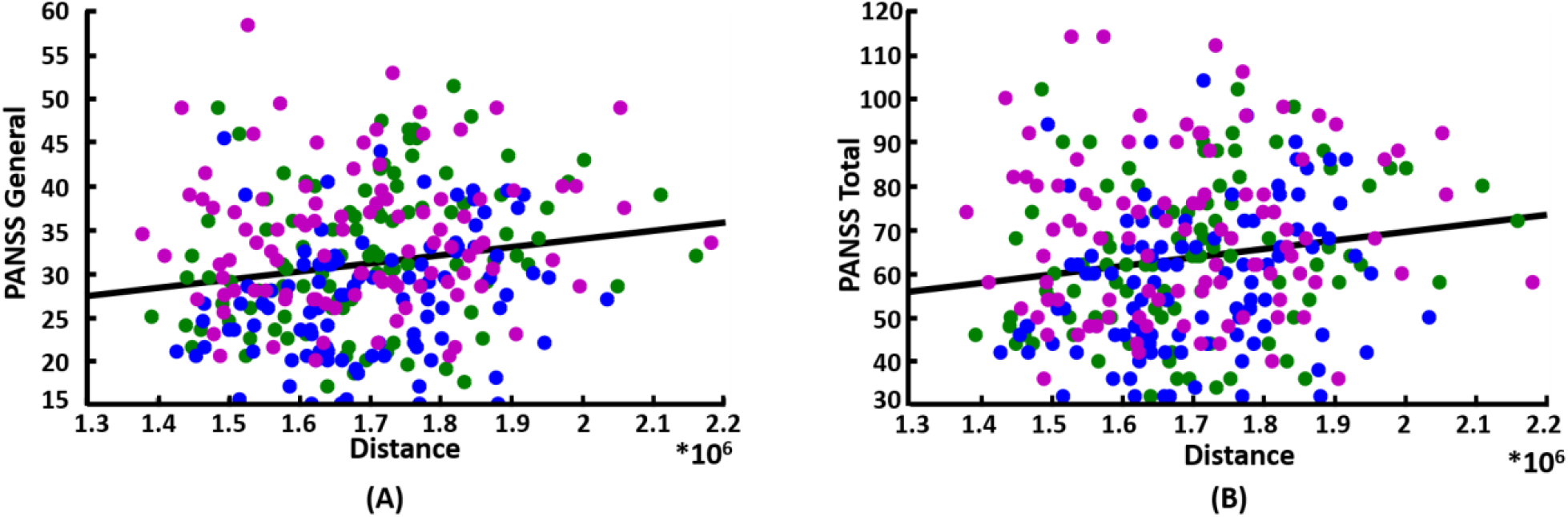
shows correlation analysis for main disease, where all diseases are combined together. The green dots are related to SZP values, blue dots to BPP, and purple to SADP. Panel A shows that distance of combined diseases can significantly predict PANSS General scores (*r*= 0.159, *p*=0.009). Panel B the same as panel A but for PANSS Total scores (*r*=0.143, *p*=0.020).

### 3.5. Comparison of network-based dynamic and static functional network connectivity

Next, we compared the state-specific FNC and static FNC for each subject across groups. We first estimated the state-specific FNC by averaging all FNCs belong a participant in each state. With three states, we have three state-specific FNC for each subject (Figure 6A step1). Then, we averaged all within and between-network functional connectivity (Figure 6A step2). For example, we have 136 connectivity features from 17 regions in CCN. Then, we averaged all 136 features to estimate inter-network functional connectivity. As such, with seven networks, we have 21 averaged within-and between-network connectivity. We calculated these 21 averaged functional connectivity features for both sFNC and state-specific dFNC features for all participants.

**Figure 6:**
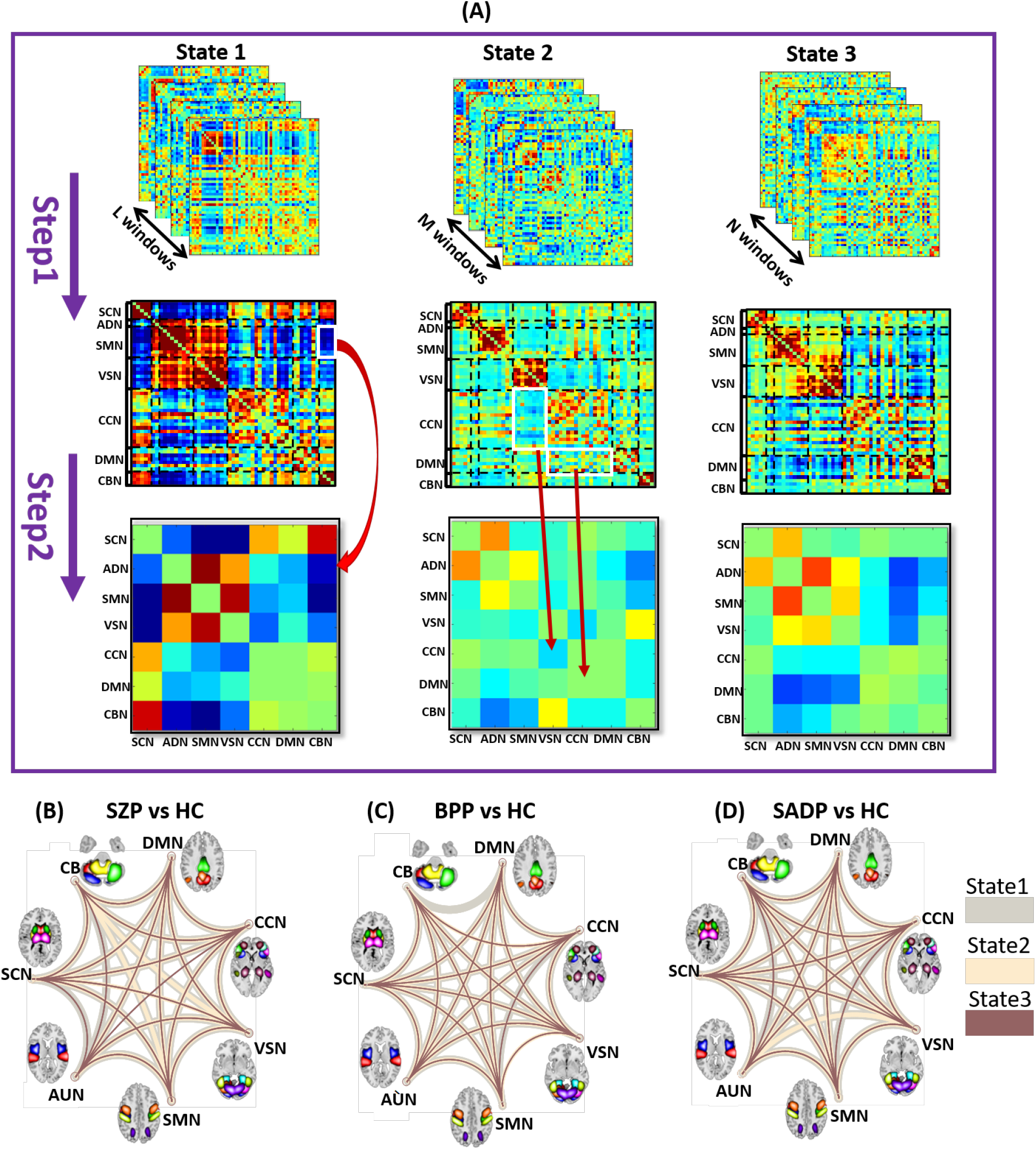
Panel A shows the procedure of calculating inter-region connectivity for one sample subject. The first row shows the number of windows that are activated in each state (L, M, and N number of windows for stat1, state2, and state3, respectively). In step1 we averaged across all the windows of each state and it resulted in second row which shows the averaged windows per state. In step2, we calculated inter-region connectivity, which is the average of all connectivity values within and between each region. Red arrows show some examples for the way we calculated inter-region connectivity from white regions in second row to specific cells in third row. Panel B to panel D are related to dynamic functional connectivity comparison in three different states of SZP vs HC, BPP vs HC, and SADP vs HC, respectively. Gray, Navajo white, and brown colored lines show significant different between regions of state1, state2, and state3, respectively. Existence of line between two regions mean there is a significant difference between those regions.

Next, we tested whether calculated features in different combination of disorders(e.g., SZP, SZR, and HC) is significant using one-way ANOVA. To summarize the results, we show figures comparing patients versus HC for each disorder (Figure 6). Figure 6B to Figure 6D compare the estimated feature for SZP v.s. HC, BPP v.s. HC, and SADP v.s. HC respectively. In this Figure 6A, the SCN/VSN (the connectivity between SCN and VSN) and ADN/CCN in state 1, the ADN/CCN and ADN/CBN in state 2, and SMN/CBN and VSN/CBN state 3 are **NOT** significant, and the rest combination of regions are significantly different (corrected *p*<0.0001). Similar results are observed in BPP vs. HC and SADP vs. HC. In addition, the results of sFNC comparison between patient vs relative group and relative vs HC group scan be seen in Figure S1. The statistical analysis shows no significant differences between any comparison of groups and regions.

## 4. Discussion

In this study, we used rs-fMRI data from 730 participants, including patients with schizophrenia (SZP), bipolar (BPP), schizoaffective disorder (SARP), their relatives (SZR, BPR, SADR), and HC group. To find overlaps among these psychotic disorders, we calculated dFNC of participants and clustered the combined dFNC of all disorders into three identical states. Then, we extracted various dFNC features from extracted state vectors of each participant and compared the statistical difference of the features in two ways: 1) within disorder (i.e., SZP/SZR/HC, BPP/BPR/HC, SADP/SADR/HC) and 2) across disorders (i.e., SZP/BPP/SADP).

This study is one of only a few studies that evaluate the dynamic functional connectivity of SAD individually and compares its overlap with SZ and BP disorders. SAD is mainly considered a sub-part of SZ, and previous studies have analyzed it combined with SZ patients in a group as a single disorder (Baker et al., 2019; Mothersill et al., 2021; Tso et al., 2021; Walther et al., 2021). Therefore, it is worth considering it a separate disorder and exploring its overlap with similar psychotic disorders. Previously, Du et al. compared whole-brain dynamic connectivity of SAD, SZ, and BP and reported that BP resembled HC in frontal connectivity, whereas SAD and SZ were more similar than other disorders (Du et al., 2017). Another study denoted that SZ, BP, and SAD show a similar pattern as measured by the prevalent functional connectivity state, highlighting that SAD is independent of SZ and BP (Du et al., 2015a). Another study by Du et al. show the ability to classify these three groups with an accuracy of over 80% (Du et al., 2020b). In line De eti al., we found an overlap between SAD and SZ disorder (Du et al., 2017). Figure 3C shows that where the OCR of SADP is significantly higher than HC in state 2, and it is replicated in state 2 of SZ disorder, where the OCR of SZP is significantly higher than HC group (Figure 3A). In addition, results of disorder comparison show a significant difference between SADP and BPP in states 3, highlighting that SAD can be viewed as an independent disorder, in line with (Du et al., 2015a). Moreover, the comparison of dFNC of the patient with HC group among different regions (Figure 6B to Figure 6D) shows many inter-region significant differences, implying that the pattern of significant connectivities has overlap in SZP, BPP, and SAD disorders, in line with other studies.

Besides the studies that highlight overlap among SZ, BP, and SAD disorders (Tamminga et al., 2013, 2014; Clementz et al., 2020; Yan et al., 2021), there is a group of studies confirming previous findings and adding the fact that the existing overlap pattern is more similar between SZ and SAD than BP. In other words, the BP activity pattern has overlap but is less identical to SZ and SAD (Ivleva et al., 2013; Amann et al., 2016; Birur et al., 2017). Even though in the patient comparison, SZP has significantly higher OCR than BPP in state2 and significantly lower OCR than BPP in state 3 (Figure 3D), our results show that the BP and SZ complement each other. As shown in Figure 3A, the HC group tends to stay in state 2 significantly lower than SZP. State 2 shows relatively lower within SMN, VSN, CCN, and DMN connectivity than the other states (Figure 2). On the other hand, Figure 3B suggests that the HC group spends more time in state 1 than BPP. State 1 shows higher connectivity within SMN, VSN, CCN, and DMN compared with other states. Therefore, aligned with previous studies, combining the findings within SZ, within BP, and between patient comparisons, we can say that although BP and SZ have not similar patterns, they are complementary.

This study is the first attempt to investigate the dynamic functional connectivity of relatives (i.e., SZR, BPR, and SADR) using rs-fMRI. Previous studies investigated relatives with patients based on behavioral scores. Hager et al. used the brief assessment of cognition in schizophrenia (BACS) score and two other behavioral scores of relatives in bipolar vs schizophrenia, and examined the clinical high risk of patients with relatives (Hager et al., 2017). They denoted that cognitive biomarkers might be crucial as indications of psychosis vulnerability. Massa et al. recorded eye-blink reaction time to a startle stimulus of SZR, BPR, and SADR patients (Massa et al., 2020). They reported that SZR has slower latency than BPR and SZR. Hochberger et al. used the BACS score of participants suffering from SZ, BP, and SAD and their relative and the control group (Hochberger et al., 2016). This work denoted that the BACS score is rooted in a similar cognitive construct in all the patients and their relatives. Even though other groups of studies used brain imaging data of relatives and compared them with patients and HCs, they did not consider the dynamic nature of brain activity (Wang et al., 2015; Hager et al., 2017; Del Fabro et al., 2021). Johnsen et al. reviewed fMRI studies which compared task-related brain alterations of SZR and BPR with SZP and BPP (Johnsen et al., 2020). They stated that the BPR group showed more alteration than the other groups in the emotion-related taks. Gromann et al using a trust game as a task, hypothesized that people having relative disorders shows lower trust behaviorally and less brain activity while doing the task (Gromann et al., 2014). Their results showed that the relative group’s behavior did not differ significantly from the controls, but they had lower activity of the right caudate and the left insula during the task. Konchel et al. evaluated differences between SZP and SZR groups in Heschl’s gyrus (HG), measuring functional connectivity between bilateral HG and the whole brain (Oertel-Knöchel et al., 2014). They showed that functional asymmetry of SZR and SZP in HG, auditory network, and connection between temporal-limbic is significantly different from HC group. In the current study, we investigated the features extracted from dFNC of the relative and patient group and compared them both within and between disorders. The results of OCR feature in within SZ comparison shows significant difference between SZP and SZR (SZP>SZR) in state 2 in which the inter-region connectivity is the lowest among all other states (Figure 3A). In comparing within BP and within SAD, we did not see any significant difference of OCR feature between relative and HC groups. The between relative disorder (Figure 3E) did not show any significant difference as well. In addition, in the other features there is not any significant differences in neither between nor within disorder comparison. The inter-region connectivity comparison of relative group with HC and with patient group shows an overlapped pattern of significant differences (Figure S1). As an example, the significant pattern for SZR-SZP shows a lot of inter region overlaps with SZR-HC significance pattern (This is the same for BP and SAD disorder).

This study used FNC to evaluate inter-region connectivity from both dynamic and static perspectives. The dFNC provides information on time-varying connectivity variations throughout time (Allen et al., 2014). Unlike static functional network connectivity (sFNC), dFNC captures the local connectivity of each window rather than providing mean connectivity (Saha et al., 2021). Recent studies have shown that rs-fMRI brain functional connectivity is hugely dynamic and can reveal the underlying mechanisms of brain connection disparities in psychotic disorders (Garrity et al., 2007; Damaraju et al., 2014; Miller et al., 2016; Zhi et al., 2018; Dong et al., 2019; Sun et al., 2019; Rey et al., 2021; Tang et al., 2022). To the best of our knowledge, there are a limited amount of studies that evaluate dFNC and sFNC simultaneously. Sendi et al. predicted the symptomatic progression of Alzheimer’s disorder (AD) using both sFNC and dFNC within the CCN region (Sendi et al., 2021e). They proved that dFNC could capture decreased connectivity between the inferior parietal lobule and the rest of CCN regions, while sFNC cannot. Saha et al. studied the dynamic versus static brain connectivity on rs-fMRI data and reported that the classification accuracy is maximum when they use either dFNC or combined dFNC and sFNC data implying that dFNC provides more information to classify HC vs. patient groups (Saha et al., 2021). In the current study, we first estimated whole-brain sFNC and then averaged the resulting connectivities within the networks to capture network-specific sFNC of each subject [Figure 1B and Figure 6A]. Then, we implement inter-network comparison using both sFNC and dFNC. As shown in Figure 6B to Figure 6D, dFNC captures the main significant difference between various regions (also in different states). However, inter-network comparison using sFNC provides no significant differences across regions. These results show that in line with previous studies, dFNC can highlight information that sFNC cannot. It is worthy of mentioning that this information is vital to study the overlap between psychotic disorders, as elaborated above.

The current study linked the extracted features from rs-fMRI od SZ, BP, and SAD with clinical scores using partial correlation analysis. Many studies have found a link between basic clinical parameters and psychotic traits (mainly focused on SZ disorder) (Hudgens-Haney et al., 2020; Bian et al., 2021; Chu et al., 2021; Huo et al., 2021; Pelizza et al., 2021; Targum et al., 2021; Teetharatkul et al., 2021). These studies are based on group-level data rather than individual patient data (Ozomaro et al., 2013). Therefore, it is essential to define new metrics and link them to the clinical scores. Choosing a metric among the various MRI metrics, on the other hand, is not easy since they focus on non-overlapping elements of brain functionality (Leaver et al., 2018). Despite studies that use fMRI metrics in clinical scores, they mainly focus on static brain connections (Leaver et al., 2018; Wang et al., 2018; Zhang et al., 2018; Ye et al., 2021). (Rashid et al., 2014). Using dFNC features, our previous study introduced a biomarker called OCR, extracted from dFNC activity of rs-fMRI of MDD patients under electroconvulsive therapy (ECT) (Dini et al., 2021). This biomarker could effectively predict the effectiveness of ECT before performing it using the data from the DMN and CCN. The current study used features extracted from dFNC of the patient group and was able to find associations between extracted features and clinical scores, using individual disorder data (Figure 4) and combined disorder features (Figure 5). The OCR feature of BP and SZ had a negative and positive correlation with general PANSS, respectively. In addition, we were able to predict total PANSS and general PANSS using the distance feature of SZ and BP. These findings align with Du et al. (Du et al., 2017), which showed the ability to predict PANSS scores using hypo/hyper-connectivity features. In addition, to test if this predictability is disorder-specific, we combined the extracted features from all disorders (Figure 5) and predicted PANSS Total and PANSS General using distance feature. Therefore, not only single disorders can predict clinical scores, but also the combined features are also able to do.

## 5. Limitation

Here we address the limitations we had in the present study. The recorded clinical scores were relatively noisy, meaning that some of the scores for some participants were not either recorded or available. We considered this issue in our analysis by removing the corresponding features and running the methods on the rest. In addition, the data was collected from multiple sites and with different TRs, that we considered them in our analysis. Also, different clinicians have recorded the scores which might add some noises to the recordings. Finally, the psychiatric classification was based on DSM-V rather than international classification of diseases (ICD).

## 6. Conclusion

This study evaluated dynamic functional network connectivity of whole brain, using rs-fMRI data of three disorders (SZ, BP, and SAD), their relatives and HC group. We clustered the brain dFNC to three identical states and found there is an overlap across these disorders using OCR feature extracted from dFNC. This is study is one of the scant studies evaluating brain dynamics of patients suffering from SAD separate from SZ disorder, highlighting its overlap with SZ group. In addition, this study is the first attempt to investigate the dynamic functional connectivity of relatives using rs-fMRI, suggesting that there is a significant difference between OCR of SZ patients and their relatives. Focusing on OCR feature extracted from dFNC, we found that the activity of BP is complementing the activity of SZ, where the OCR of HC group is higher than BP in a state that the connectivity of nodes is lowest among the other states, while the OCR of HC is lower than SZ in a state which shows the highest connectivity features. In addition, we compared the dFNC activity vs sFNC results which showed that dFNC captures the information that are ignored by sFNC, suggesting a highly informative method. Finally, we could significantly find associations between OCR/distance features and clinical scores. We found that the more time BP groups spend in state 2 the less PANSS general score is reported, while the more time SZ patients spend in state 1 the higher PANSS general score is reported. We also could significantly find associations between distance feature and clinical scores of SZ and BP groups. In brief, this study provides a focus on functional connectivity dynamics of whole brain network of three psychiatric disorders and highlights the overlap across them introducing group discriminative neurophysiological features, which are associated with the PANSS scores.

## 7. Data availability statement

The raw data supporting the conclusions of this article will be made available by the authors, without undue reservation.

## 8. Author Contributions

HD and MS developed the study, conducted data analysis, interpreted the results, and wrote the original manuscript draft. LB and TR provided a critical review of the initial draft, and edited the original draft. VC developed the study, interpreted the results, edited the original draft, and provided critical review to the initial draft. All authors approved the final manuscript.

## 10. Acknowledgments

We thank those who helped collect these valuable data, and thanks to our funding institutions NIH and NFS.

