## Supplemental Information for "Overlap across psychotic disorders: A functional network connectivity analysis"

** In all tables the significant differences are highlighted in yellow, and all of the p-values are corrected as explained in the paper.

| Table S1: Results of between and within comparison using HMM feature | | | | | | | | | | |
| --- | --- | --- | --- | --- | --- | --- | --- | --- | --- | --- |
|  |  | 1-1 | 1-2 | 1-3 | 2-1 | 2-2 | 2-3 | 3-1 | 3-2 | 3-3 |
| Within  SZ | SZP vs SZR | 0.085 | 0.850 | 0.782 | 0.864 | 0.517 | 0.622 | 0.623 | 0.935 | 0.323 |
|  | SZP vs HC | 0.019 | 0.789 | 0.992 | 0.207 | 0.287 | 0.157 | 0.016 | 0.812 | 0.236 |
|  | SZR vs HC | 0.726 | 0.969 | 0.894 | 0.309 | 0.368 | 0.579 | 0.232 | 0.601 | 0.939 |
| Within  BP | BPP vs BPR | 0.897 | 0.310 | 0.881 | 0.821 | 0.978 | 0.278 | 0.714 | 0.908 | 0.681 |
|  | BPP vs HC | 0.455 | 0.314 | 0.886 | 0.115 | 0.322 | 0.605 | 0.995 | 0.827 | 0.979 |
|  | BPR vs HC | 0.731 | 0.999 | 0.999 | 0.345 | 0.430 | 0.834 | 0.658 | 0.984 | 0.797 |
| Within  SAD | SADP vs SADR | 0.987 | 0.982 | 0.943 | 0.931 | 0.369 | 0.286 | 0.935 | 0.970 | 0.837 |
|  | SADP vs HC | 0.380 | 0.955 | 0.944 | 0.186 | 0.020 | 0.295 | 0.978 | 0.984 | 0.798 |
|  | SADR vs HC | 0.380 | 0.915 | 0.999 | 0.209 | 0.316 | 0.194 | 0.815 | 0.872 | 0.976 |
| Between  Main | SZP vs BPP | 0.451 | 0.922 | 0.958 | 0.823 | 0.499 | 0.149 | 0.502 | 0.999 | 0.090 |
|  | SZP vs SADP | 0.575 | 0.999 | 0.874 | 0.991 | 0.858 | 0.776 | 0.734 | 0.711 | 0.494 |
|  | BPP vs SADP | 0.852 | 0.936 | 0.720 | 0.753 | 0.210 | 0.416 | 0.965 | 0.735 | 0.598 |
| Between  Relative | SZR vs BPR | 0.999 | 0.959 | 0.545 | 0.819 | 0.992 | 0.283 | 0.503 | 0.757 | 0.771 |
|  | SZR vs SADR | 0.890 | 0.914 | 0.439 | 0.899 | 0.969 | 0.957 | 0.966 | 0.947 | 0.986 |
|  | BPR vs SADR | 0.889 | 0.877 | 0.999 | 0.852 | 0.933 | 0.441 | 0.324 | 0.915 | 0.869 |

| Table S2: Results of between and within comparison using OCR feature | | | | |
| --- | --- | --- | --- | --- |
|  |  | State 1 | State 2 | State 3 |
| Within  SZ | SZP vs SZR | 0.109 | 0.021 | 0.509 |
|  | SZP vs HC | 0.068 | 0.000 | 0.072 |
|  | SZR vs HC | 0.896 | 0.218 | 0.431 |
| Within  BP | BPP vs BPR | 0.264 | 0.995 | 0.992 |
|  | BPP vs HC | 0.045 | 0.454 | 0.976 |
|  | BPR vs HC | 0.579 | 0.981 | 0.989 |
| Within  SAD | SADP vs SADR | 0.790 | 0.108 | 0.250 |
|  | SADP vs HC | 0.109 | 0.001 | 0.348 |
|  | SADR vs HC | 0.249 | 0.225 | 0.982 |
| Between  Main | SZP vs BPP | 0.998 | 0.012 | 0.004 |
|  | SZP vs SADP | 0.865 | 0.658 | 0.818 |
|  | BPP vs SADP | 0.905 | 0.080 | 0.017 |
| Between  Relative | SZR vs BPR | 0.87 | 0.838 | 0.238 |
|  | SZR vs SADR | 0.758 | 0.992 | 0.593 |
|  | BPR vs SADR | 0.987 | 0.503 | 0.669 |

| Table S3: Results of between and within comparison using Transition to one state feature | | | | |
| --- | --- | --- | --- | --- |
|  |  | State 1 | State 2 | State 3 |
| Within  SZ | SZP vs SZR | 0.291 | 0.157 | 0.578 |
|  | SZP vs HC | 0.029 | 0.238 | 0.075 |
|  | SZR vs HC | 0.659 | 0.995 | 0.376 |
| Within  BP | BPP vs BPR | 0.486 | 0.608 | 0.478 |
|  | BPP vs HC | 0.353 | 0.869 | 0.832 |
|  | BPR vs HC | 0.978 | 0.817 | 0.709 |
| Within  SAD | SADP vs SADR | 0.844 | 0.537 | 0.521 |
|  | SADP vs HC | 0.420 | 0.543 | 0.335 |
|  | SADR vs HC | 0.657 | 0.908 | 0.991 |
| Between  Main | SZP vs BPP | 0.842 | 0.795 | 0.299 |
|  | SZP vs SADP | 0.983 | 0.832 | 0.709 |
|  | BPP vs SADP | 0.992 | 0.801 | 0.647 |
| Between  Relative | SZR vs BPR | 0.812 | 0.811 | 0.101 |
|  | SZR vs SADR | 0.998 | 0.875 | 0.467 |
|  | BPR vs SADR | 0.835 | 0.979 | 0.558 |

| Table S4: Results of between and within comparison using All Transitions and Distance features | | | |
| --- | --- | --- | --- |
|  |  | All Transitions | Distance |
| Within  SZ | SZP vs SZR | 0.093 | 0.889 |
|  | SZP vs HC | 0.003 | 0.754 |
|  | SZR vs HC | 0.408 | 0. 664 |
| Within  BP | BPP vs BPR | 0.255 | 0.942 |
|  | BPP vs HC | 0.507 | 0.838 |
|  | BPR vs HC | 0.754 | 0.969 |
| Within  SAD | SADP vs SADR | 0.097 | 0.736 |
|  | SADP vs HC | 0.086 | 0.691 |
|  | SADR vs HC | 0.949 | 0.434 |
| Between  Main | SZP vs BPP | 0.207 | 0.979 |
|  | SZP vs SADP | 0.687 | 0.890 |
|  | BPP vs SADP | 0.564 | 0.792 |
| Between  Relative | SZR vs BPR | 0.297 | 0.634 |
|  | SZR vs SADR | 0.818 | 0.926 |
|  | BPR vs SADR | 0.570 | 0.703 |

CORRELATION

| Table S5: Results of correlation analysis using OCR feature | | | | |
| --- | --- | --- | --- | --- |
|  |  | State 1 | State 2 | State 3 |
| SZP | PANSS Positive | P=0.442  R=0.154 | P=0.530  R=-0.098 | P=0.778  R=-0.031 |
|  | PANSS Negative | P=0.142  R=0.177 | P=0.258  R=-0.182 | P=0.678  R=0.044 |
|  | PANSS Gentotal | P=0.420  R=0.115 | P=0.580  R=-0.138 | P=0.638  R=0.050 |
|  | PANSS Total | P=0.313  R=0.173 | P=0.180  R=-0.165 | P=0.793  R=0.028 |
| BPP | PANSS Positive | P=0.132  R=0.222 | P=0.583  R=-0.096 | P=0. 691  R=-0.044 |
|  | PANSS Negative | P=0.6720  R=-0.046 | P=0.083  R=-0.214 | P=0. 071  R=0.246 |
|  | PANSS Gentotal | P=0.142  R=0.185 | P=0.043  R=-0.269 | P=0.161  R=0.153 |
|  | PANSS Total | P=0.203  R=0.166 | P=0.073  R=-0.249 | P=0.191  R=0.145 |
| SADP | PANSS Positive | P=0.774  R=0.031 | P= 0.776  R=-0.031 | P=0.915  R=0.011 |
|  | PANSS Negative | P=0.823  R=0.024 | P=0.897  R=-0.076 | P=0.981  R=0.067 |
|  | PANSS Gentotal | P=0.843  R=0.015 | P=0.897  R=-0.067 | P=0.981  R=0.064 |
|  | PANSS Total | P=0.773  R=0.031 | P=0.973  R=-0.072 | P=0.991  R=0.056 |

|  | Table S6: Results of correlation analysis using HMM feature | | | | | | | | | |
| --- | --- | --- | --- | --- | --- | --- | --- | --- | --- | --- |
|  |  | 1-1 | 1-2 | 1-3 | 2-1 | 2-2 | 2-3 | 3-1 | 3-2 | 3-3 |
| SZP | PANSS Positive | P=0.715  R=0.039 | P=0.364  R=-0.097 | P=0.599  R=0.056 | P=0.314  R=0.107 | P=0.586  R=-0.058 | P=0.630  R=0.051 | P=0.835  R=0.022 | P=0.923  R= -0.010 | P=0.108  R=-0.171 |
|  | PANSS Negative | P=0.492  R=0.181 | P=0.900  R=-0.013 | P=0.280  R=0.115 | P= 0.186  R=0.141 | P=0.799  R=-0.027 | P=0.312  R=0.108 | P=0. 116  R= 0.266 | P=0.420  R=-0.086 | P=0.973  R=-0.003 |
|  | PANSS Gentotal | P=0.397  R=0.090 | P=0.796  R=-0.027 | P=0.493  R=0.073 | P=0.104  R=0.173 | P=0.614  R=0.054 | P=0.819  R=0.024 | P=0.706  R=0.040 | P=0.975  R=0.003 | P=0.786  R=-0.029 |
|  | PANSS Total | P=0.257  R=0.121 | P=0.622  R=-0.052 | P=0.369  R=0.096 | P=0.108  R=0.171 | P=0.963  R=-0.004 | P=0.523  R=0.068 | P=0.257  R=0.121 | P=0.761  R=-0.032 | P=0.474  R=-0.076 |
| BPP | PANSS Positive | P=0.593  R=0.059 | P=0.206  R=-0.141 | P=0.167  R=-0.153 | P=0.155  R=0.158 | P=0.539  R=-0.068 | P=0.380  R=-0.098 | P=0.168  R=0.153 | P=0.710  R=-0.041 | P=0.970  R=0.004 |
|  | PANSS Negative | P=0.649  R=-0.051 | P=0.309  R=-0.113 | P=0.284  R=-0.119 | P=0.557  R=0.065 | P=0.598  R=-0.059 | P=0.564  R=0.064 | P=0.843  R=-0.022 | P=0.834  R=0.023 | P=0.343  R=0.223 |
|  | PANSS Gentotal | P=0.776  R=0.031 | P=0.194  R=-0.144 | P=0.460  R=-0.082 | P=0.477  R=0.220 | P=0.534  R=-0.069 | P=0.856  R=0.020 | P=0.698  R=0.043 | P=0.995  R=-0.000 | P=0.292  R=0.117 |
|  | PANSS Total | P=0.836  R=0.023 | P=0.142  R=-0.163 | P=0.232  R=-0.133 | P=0.747  R=0.198 | P=0.474  R=-0.080 | P=0.987  R=-0.001 | P=0.536  R=0.069 | P=0.948  R=-0.007 | P=0.227  R=0.134 |
| SADP | PANSS Positive | P=0.593  R=0.059 | P=0.206  R=-0.140 | P=0.167  R=-0.153 | P=0.155  R=0.158 | P=0.539  R=-0.067 | P=0.380  R=-0.098 | P=0.168  R=0.153 | P=0.710  R=-0.041 | P=0.970  R=0.004 |
|  | PANSS Negative | P=0.649  R=-0.051 | P=0.309  R=-0.113 | P=0.284  R=-0.119 | P=0.557  R=0.065 | P=0.598  R=-0.059 | P=0.564  R=0.064 | P=0.843  R=-0.022 | P=0.834  R=0.023 | P=0.436  R=0.223 |
|  | PANSS Gentotal | P=0.776  R=0.031 | P=0.194  R=-0.144 | P=0.460  R=-0.082 | P=0.247  R=0.220 | P=0.534  R=-0.069 | P=0.856  R=0.020 | P=0.698  R=0.043 | P=0.995  R=-0.006 | P=0.292  R=0.117 |
|  | PANSS Total | P=0.653  R= 0.049 | P=0.882  R=0.016 | P=0.411  R=0.090 | P=0.746  R=-0.038 | P=0.471  R=0.079 | P=0.574  R=-0.062 | P=0.399  R=-0.093 | P=0.994  R=0.000 | P=0.849  R=0.021 |

| Table S7: Results of correlation analysis using Transition to a state feature | | | | |
| --- | --- | --- | --- | --- |
|  |  | State 1 | State 2 | State 3 |
| SZP | PANSS Positive | P=256  R=0.121 | P=0.131  R=-0.214 | P=0.279  R=-0.143 |
|  | PANSS Negative | P=0.034  R=0.261 | P=0.145  R=-0.177 | P=0.726  R=-0.037 |
|  | PANSS Gentotal | P=0.255  R=0.1836 | P=0.324  R=-0.1324 | P=0.597  R=-0.0567 |
|  | PANSS Total | P=0.106  R=0.223 | P=0.085  R=-0.2026 | P=0.392  R=-0.0918 |
| BPP | PANSS Positive | P=0.4077  R=0.092 | P=0.5087  R=-0.0740 | P=0.3738  R=-0.0995 |
|  | PANSS Negative | P=0.2786  R=-0.1211 | P=0.4467  R=-0.0852 | P=0.9012  R=0.0139 |
|  | PANSS Gentotal | P=0.7022  R=0.0429 | P=0.6130  R=-0.0567 | P=0.8595  R=0.0199 |
|  | PANSS Total | P=0.8596  R=0.0198 | P=0.4638  R=-0.0820 | P=0.8774  R=-0.0173 |
| SADP | PANSS Positive | P=0.6981  R=0.0429 | P=0.8410  R=0.0222 | P=0.7997  R=0.0281 |
|  | PANSS Negative | P=0.9153  R=-0.0118 | P=0.6138  R=0.0559 | P=0.6043  R=0.0574 |
|  | PANSS Gentotal | P=0.6750  R=-0.0467 | P=0.6455  R=-0.0512 | P=0.9696  R=-0.0043 |
|  | PANSS Total | P=0.9371  R=-0.0088 | P=0.9138  R=-0.0121 | P=0.8584  R=0.0199 |

| Table S8: Results of correlation analysis using All Transition and Distance to a state features | | | |
| --- | --- | --- | --- |
|  |  | All Transitions | Distance |
| SZP | PANSS Positive | P=0.2657  R=-0.1193 | P=0.064  R=0.1967 |
|  | PANSS Negative | P=0.995  R=-0.0006 | P=0.283  R=0.1150 |
|  | PANSS Gentotal | P=0.859  R=-0.0190 | P=0.015  R=0.2571 |
|  | PANSS Total | P=0.627  R=-0.0521 | P=0.0275  R= 0.2338 |
| BPP | PANSS Positive | P=0.630  R=-0.0539 | P=0.589  R=0.0604 |
|  | PANSS Negative | P=0.440  R=-0.0864 | P=0.084  R=0.1915 |
|  | PANSS Gentotal | P=0.988  R=0.0016 | P=0.027  R=0.2438 |
|  | PANSS Total | P=0.707  R= -0.0421 | P=0.050  R=0.2165 |
| SADP | PANSS Positive | P=0.704  R=0.0420 | P=0.788  R=0.0297 |
|  | PANSS Negative | P=0.660  R=0.0486 | P=0.729  R=0.0383 |
|  | PANSS Gentotal | P=0.698  R=-0.0432 | P=0.484  R=0.0778 |
|  | PANSS Total | P=0.988  R=0.0016 | P=0.524  R=0.0709 |

Grand Correlation

| Table S9: Results of correlation analysis using combined OCR features | | | |
| --- | --- | --- | --- |
|  | State1 | State2 | State3 |
| PANSS Positive | P=0.057  R=0.1444 | P=0.765  R=-0.0185 | P=0.222  R=-0.0894 |
| PANSS Negative | P=0.281  R=0.0813 | P=0.317  R=-0.0999 | P=0.450  R=0.0467 |
| PANSS Gentotal | P=0.224  R=0.1102 | P=0.117  R=-0.1086 | P=0.582  R=0.0341 |
| PANSS Total | P=0.091  R=0.1337 | P=0.186  R=-0.0951 | P=0.977  R=0.0018 |

|  | Table S10: Results of correlation analysis using combined HMM features | | | | | | | | |
| --- | --- | --- | --- | --- | --- | --- | --- | --- | --- |
|  | 1-1 | 1-2 | 1-3 | 2-1 | 2-2 | 2-3 | 3-1 | 3-2 | 3-3 |
| PANSS Positive | P=0.5075  R=0.0410 | P=0.3845  R=-0.1066 | P=0.6431  R=-0.0287 | P=0.4557  R=0.0703 | P=0.9040  R=-0.0075 | P=0.2339  R=-0.0736 | P=0.7466  R=0.0200 | P=0.7905  R=0.0165 | P=0.5070  R=-0.0781 |
| PANSS Negative | P=0.7548  R=0.0193 | P=0.1890  R=-0.0813 | P=0.6979  R=-0.0240 | P=0.4580  R=0.0460 | P=0.8001  R=-0.0157 | P=0.4666  R=0.0451 | P=0.2982  R=-0.0644 | P=0.8384  R=0.0126 | P=0.5770  R=0.0345 |
| PANSS Gentotal | P=0.7345  R=-0.0210 | P=0.1622  R=-0.1154 | P=0.4455  R=-0.0473 | P=0.1407  R=0.1266 | P=0.9320  R=0.0053 | P=0.8174  R=-0.0143 | P=0.7034  R=-0.0236 | P=0.5911  R=0.0333 | P=0.9196  R=0.0063 |
| PANSS Total | P= 0.8503  R= 0.0117 | P=0.1495  R=-0.1215 | P=0.5052  R=-0.0413 | P=0.1880  R=0.1056 | P=0.9384  R=-0.0048 | P=0.7676  R=-0.0183 | P=0.6741  R=-0.0261 | P=0.6647  R=0.0269 | P=0.8118  R=-0.0148 |

| Table S11: Results of correlation analysis using combined Transition to a state features | | | |
| --- | --- | --- | --- |
|  | State1 | State2 | State3 |
| PANSS Positive | P=0.1637  R=0.0861 | P=0.1125  R=-0.0981 | P=0.1480  R=-0.0894 |
| PANSS Negative | P=0.3712  R=0.0554 | P=0.1562  R=-0.0877 | P=0.8597  R=-0.0109 |
| PANSS Gentotal | P=0.2774  R=0.0673 | P=0.2600  R=-0.0698 | P=0.7923  R=-0.0163 |
| PANSS Total | P=0.1750  R= 0.0840 | P=0.1089  R=-0.0993 | P=0.4832  R=-0.0435 |

| Table S12: Results of correlation analysis using combined All Transitions and Distance features | | |
| --- | --- | --- |
|  | All Transitions | Distance |
| PANSS Positive | P=0.370  R=-0.055 | P=0.1844  R=0.0821 |
| PANSS Negative | P=0.7219  R=-0.0221 | P=0.1225  R=0.0955 |
| PANSS Gentotal | P=0.8427  R=-0.0123 | P=0.0097  R=0.1596 |
| PANSS Total | P=0.5976  R=-0.0328 | P=0.0200  R=0.1436 |

| Table S13: Scanning parameters, subject number, and sliding window parameters of all sites. | | | | | | | | | |
| --- | --- | --- | --- | --- | --- | --- | --- | --- | --- |
| Site | TR (ms) | TE (ms) | Flip angle (degree) | Slices number | Voxel Size (mm) | Time points | Subject number | Window number | Window length (s) |
| Baltimore | 2210 | 30 | 70 | 36 | 3.4x3.4x4 | 140 | 133 (49 HCs, 23 BPP, 25 SAD and 36 SZ patients) | 115 | 44.2 |
| Boston | 3000 | 27 | 60 | 30 | 3.4x3.4x5 | 100 | 19 (9 HCs, 4 BPP, 2 SAD and 4 SZ patients) | 75 | 60.0 |
| Chicago | 1775 | 27 | 60 | 29 | 3.4x3.4x5 | 210 | 137 (52 HCs, 40 BPP, 25 SAD, 20 SZ patients) | 185 | 33.5 |
| Dallas | 1500 | 27 | 60 | 29 | 3.4x3.4x5 | 210 | 133 (57 HCs, 21 BPP, 37 SAD and 18 SZ patients) | 185 | 30 |
| Detroit(A) | 1570 | 22 | 60 | 29 | 3.4x3.4x5 | 210 | 14 (8 HCs, 3 BPP, and 3 SZ patients) | 185 | 31.4 |
| Detroit(B) | 1720 | 23 | 60 | 29 | 3.8x3.8x5 | 210 | 49 (15 HCs, 19 BPP, 5 SAD, and 10 SZ patients) | 185 | 34.4 |
| Hartford | 1500 | 27 | 70 | 29 | 3.4x3.4x5 | 210 | (48 HCs, 30 BPP, 38 SAD and 22 SZ patients) | 185 | 30 |

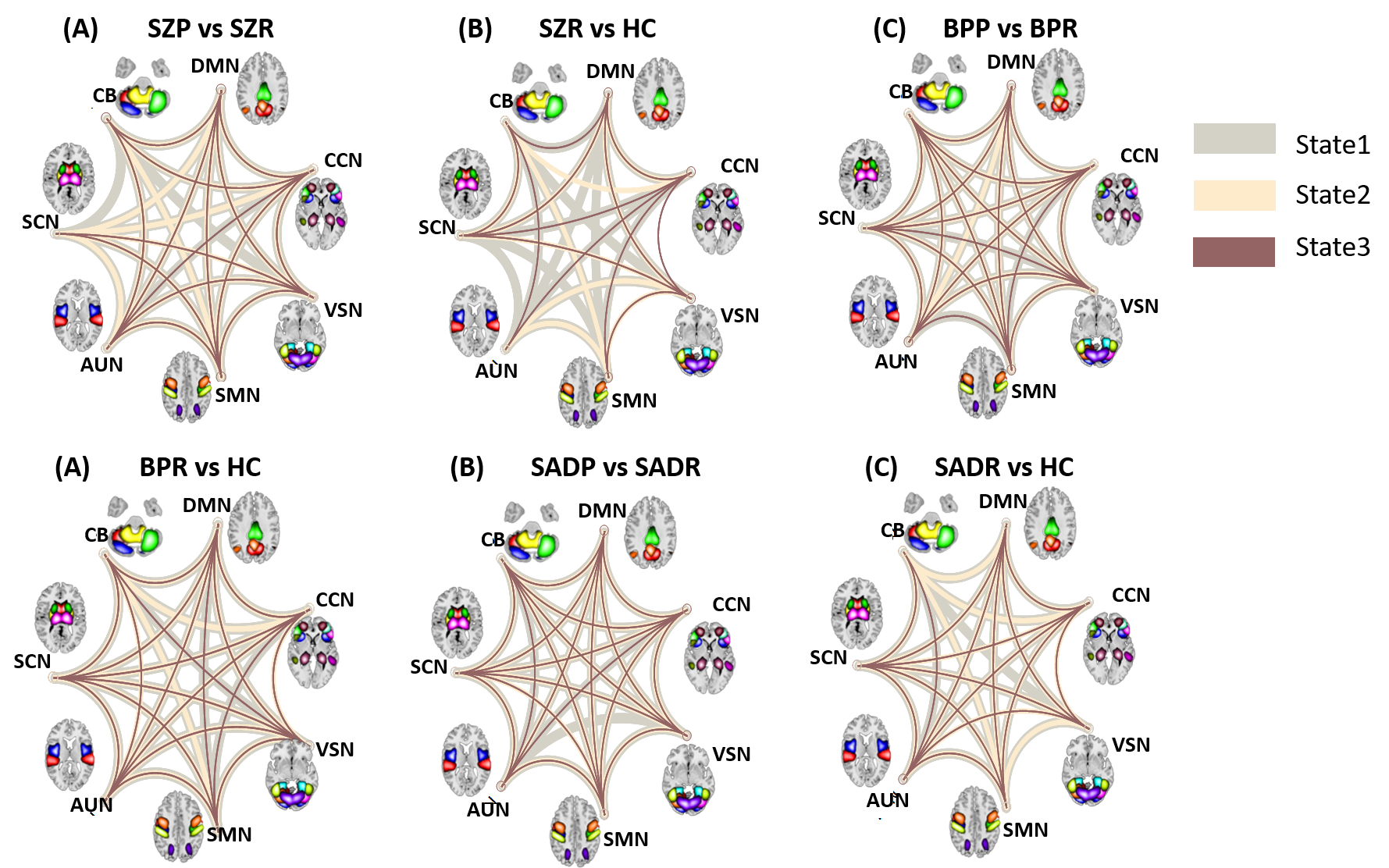

**FigureS1:** Dynamic and static functional connectivity comparison between regions for relative disease vs main disease and relative disease vs HC group. Gray, Navajo white, and brown colored lies show significant different between regions of state1, state2, and state3, respectively. Existence of line between two regions mean there is a significant difference between those regions.
